# Associative memory networks for graph-based abstraction

**DOI:** 10.1101/2020.12.24.424371

**Authors:** Tatsuya Haga, Tomoki Fukai

**Affiliations:** Okinawa Institute of Science and Technology, Onna-son, Okinawa, Japan

**Author notes:** correspondence (TH), (TF).

## Abstract

Our cognition relies on the ability of the brain to segment hierarchically structured events on multiple scales. Recent evidence suggests that the brain performs this event segmentation based on the structure of state-transition graphs behind sequential experiences. However, the underlying circuit mechanisms are only poorly understood. In this paper, we propose an extended attractor network model for the graph-based hierarchical computation, called as Laplacian associative memory. This model generates multiscale representations for communities (clusters) of associative links between memory items, and the scale is regulated by heterogenous modulation of inhibitory circuits. We analytically and numerically show that these representations correspond to graph Laplacian eigenvectors, a popular method for graph segmentation and dimensionality reduction. Finally, we demonstrate that our model with asymmetricity exhibits chunking resembling to hippocampal theta sequences. Our model connects graph theory and the attractor dynamics to provide a biologically plausible mechanism for abstraction in the brain.

## Introduction

The brain builds a hierarchical knowledge structure through the abstraction of groups and segments. This ability of the brain is essential for various cognitive functions such as chunking of items which increases the number of items kept in a limited capacity of working memory^1^, segmentation of words which is essential for learning and comprehension of language^2–4^, and temporal abstraction of repeated sequential actions which accelerates reinforcement learning^5^.

Experimental evidence suggest that the brain performs segmentation based on the graph structures behind experiences. When the brain experiences a sequence of events, it learns temporal associations between successive events and eventually captures the structure of the state-transition graph behind the experience. It has been shown that the event segmentation performed by human subjects behaviorally reflects community structures (or clusters) of such state-transition graphs and, neurobiologically, sensory events within a same community are represented by more similar activity patterns than those belonging to different communities^6,7^. Computationally, such graph segmentation of events is considered to benefit the temporal abstraction of actions in reinforcement learning^8,9^. Furthermore, graph-based representations can explain many characteristics of place cells and entorhinal grid cells^10^.

Despite their behavioral and representational evidence, the biological mechanism that creates graph-based representations remains unknown. Conventionally, the circuit-level mechanisms in hippocampal and cortical processing have been modeled as attractor-based associative memory networks^11–13^. Experiments have revealed some hallmarks of associative memory networks such as Hebbian learning (as spike-timing dependent plasticity)^14,15^, pattern completion and attractor states^16–19^ in the brain. In the context of associative memory, temporal associations between successive events can be modeled as hetero-associations between successively activated cell assemblies through Hebbian learning^20–23^. This learning scheme creates correlated attractors from originally uncorrelated stimuli. Correlations depends on the temporal distance between memorized events along the event sequence, which quantitatively agrees with neural recordings from the monkey brain^24,25^. Such correlated attractors, and hence the class of associative memory models, can be potentially extended to offer a biologically plausible representational basis for more general graphical structures. However, this hypothesis has not been examined.

In this study, we propose a generalized class of associative memory networks^20–23^ that performs graph-based segmentation and abstraction. We make two major extensions here. First, we generalize the one-dimensional sequential structure of temporal associations in the conventional model^20–23^ to arbitrary symmetric graphs. Second, we allow the model to have negative associative weights which can be interpreted as assembly-specific inhibition^26^. We found that this network generates mixed representations that are shared by multiple memory items within same communities in the graph, which fits with human experiments^6,7^. We mathematically revealed that attractors in our model are related to graph Laplacian eigenvectors, a popular mathematical method to perform graph segmentation^27,28^ and nonlinear dimensionality reduction^29^. Because of this property, we call our model as Laplacian associative memory (LAM), and demonstrate that LAM is applicable to problems related to graph Laplacian eigenvectors such as subgoal finding in reinforcement learning^8,9^. The scale of representations (the size of represented communities) is modulated by the relative strength of local and global inhibitory circuits, which indicates an active role of target-specific inhibition^26^ and inhomogeneous neuromodulation of inhibitory circuits^30,31^. Finally, we show that LAM with asymmetric weight bias generates chunked sequential activities observed in hippocampus^32,33^. To our knowledge, our model describes the first theoretical results that connect associative memory networks and graph theory, providing a biologically plausible dynamical mechanism for the hierarchical abstraction in the brain.

## Results

### Laplacian associative memory model

Laplacian associative memory (LAM) is a novel class of Hopfield-type recurrent network model^11–13,20,23^. Let us define a network of *N* binary units *x*_*i*_(*t*) (*i* = 1, …, *N*; 0 ≤ *x*_*i*_(*t*) ≤ 1) with the dynamics:

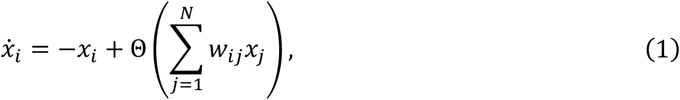

where *w*_*ij*_ are synaptic weights and Θ(*x*) is a step function (Θ(*x*) = 1 if *x* > 0, otherwise Θ(*x*) = 0). We assume that each memory item (e.g. sensory stimuli, places or events) is represented by a 0-1 binary random memory pattern 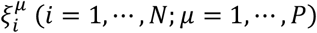 with sparsity *p* 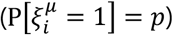. We set synaptic weights from these memory patterns as

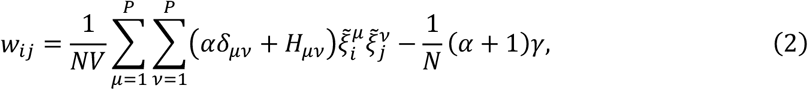

where 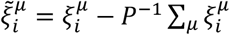 and *V* = *p*(1 − *p*). The term *αδ*_*μν*_ represents auto-association within each item, where *δ*_*μν*_ is Kronecker delta and *α* is a modifiable parameter that determines the strength of auto-association. On the other hand, *H*_*μν*_ is hetero-associative weights between memory items *μ* and *ν* (*H*_*μμ*_ = 0). The parameter *γ* ≥ 0 gives an additional global inhibitory effect^13^. In short, this network stores multiple cell assemblies (*P* memory patterns) through auto-associative Hebbian learning and further links them through hetero-associative Hebbian learning (Figure 1, left). Assuming Hebbian learning between successively activated cell assemblies^22,25^, we construct hetero-associative weights from a normalized adjacency matrix of a state-transition graph, or generally, other graphs such as semantic relationships (see Methods for the detail). Thus, the structure of hetero-associative links is a graph reflecting the statistical structure behind experiences in which we may find some communities.

LAM can be regarded as the generalization of previous associative memory models. When *α* > 0 and all *H*_*μν*_ are zero, LAM is analogous to conventional Hopfield-type model storing biased memory patterns^11–13^. If only adjacent items are associated (*H*_*μ*,*μ*+1_ = *H*_*μ*+1,*μ*_ > 0, and all other *H*_*μν*_ are zero) so that associative links form a one-dimensional chain, the model coincides with an associative memory model for a temporal sequence^20,23^. However, unlike those previous models, LAM can also take other arbitrary hetero-associative link structures. Furthermore, we do not restrict the parameter *α* to be positive, allowing inhibitory auto-association. We found unique behaviors of LAM mostly in the regime of negative auto-association, which has not been extensively investigated previously.

**Figure 1.**
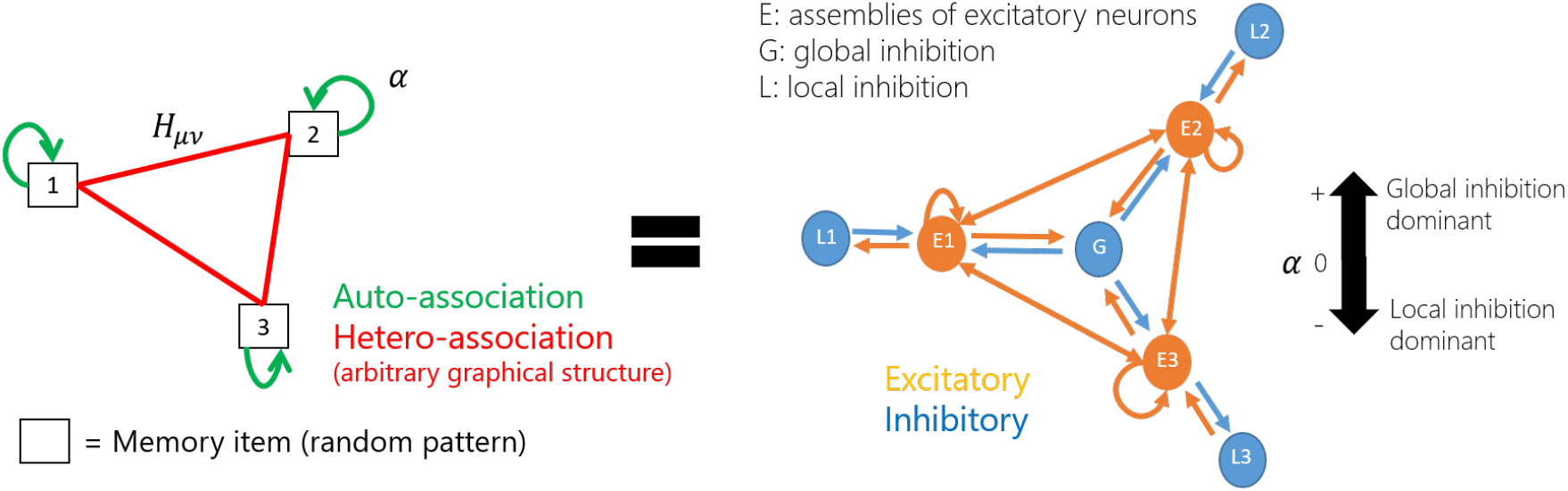
Schematics of Laplacian associative memory (LAM) model. Left: associative memory network model with auto-association and hetero-association. The parameter *α* indicates the auto-associative strength. Right: Equivalent biological neural network model which contains local (assembly-specific) and global (non-specific) inhibition. *α* indicates the ratio between local and global inhibition.

We clarify biological interpretation of the model by decomposition of excitatory and inhibitory components. As in a previous work^23^, we can decompose the weights as

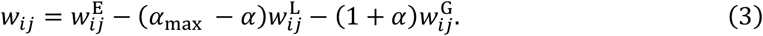

In the range −1 < *α* < *α*_max_, decomposed weights 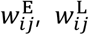, and 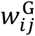 (always non-negative values) represent excitatory connections, pattern-specific local inhibition and non-selective global inhibition, respectively (see Methods for the detailed description). Therefore, LAM can be regarded as a circuit with local and global inhibition, in which the parameter *α* determines the ratio between strengths of two types of inhibitory circuits (Figure 1, right). Biologically, the difference of *α* may correspond to anatomical inhomogeneity of interneurons. Otherwise, the balance of inhibition can be changed through inhomogeneous modulation of interneurons by acetylcholine^30,31^.

### Multiscale representation of community structures in LAM

To demonstrate representions in LAM, we tested three representative graph structures. The first one is the graph used previously to study how humans segment temporal sequences obeying probabilistic state-transition rules^6,7^ (Figure 2a). The second is karate club network^34^ which is a popular dataset for testing community detection methods in graph theory (Figure 2f). The third is a graph representing the structure of compartmentalized rooms (Figure 2k) which is often used as the state-transition graph for reinfocement learning^5,8–10^. For each graph, we assigned a random binary pattern to each node and constructed a LAM network by setting hetero-associative weights *H*_*μν*_ from adjacency matrices of the graph. We initialized the activity pattern of LAM with one of memory patterns (trigger stimulus corresponding to each node), and simulated dynamics of the network for sufficiently long time to converge to an attractor state. We regard that attractor pattern as a neural representation of the node.

**Figure 2.**
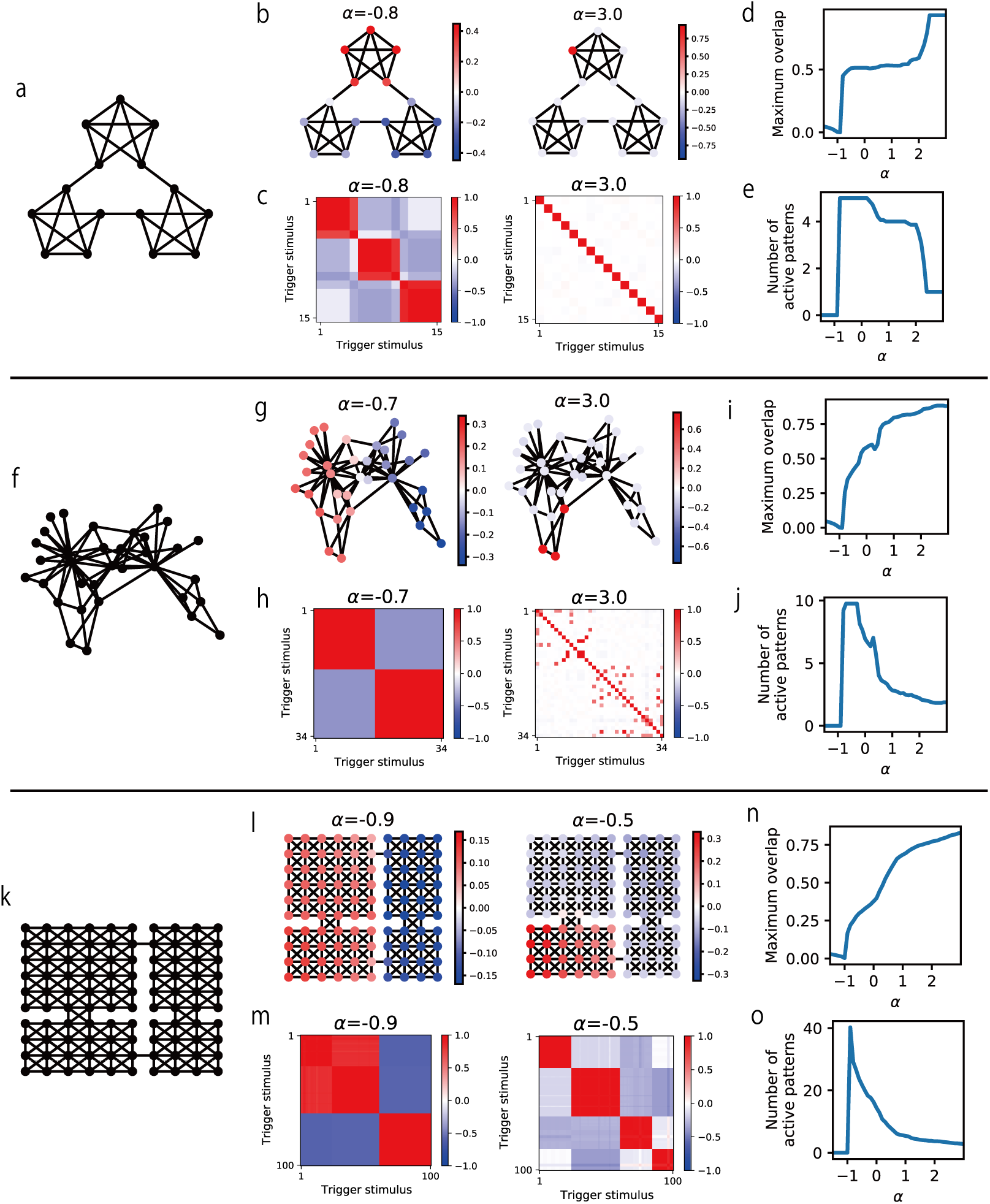
LAM generates multi-scale representations for community structures. (a) Graph used in Schapiro et al. (2013)^6^. (b) Pattern overlaps of example attractor patterns. (c) Correlation matrices between attractor patterns triggered by different initial patterns obtained. (d) Maximum pattern overlaps obtained by various values of *α*. (e) Numbers of active patterns obtained by various values of *α*. In d and e, values from all attractor patterns triggered by different initial patterns are averaged for each *α*. (f) Karate club network^34^. (g-j) Results for Karate-club network. (k) A compartmentalized room structure used in the context of reinforcement learning^5,9,10,37^ (four-room graph). (l-o) Results for four-room graph.

For each attractor state, we calculated an index called as pattern overlaps to evaluate the degree of retrieval of each memory pattern:

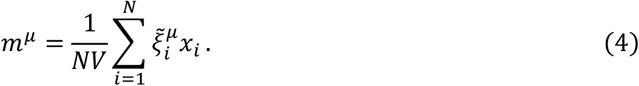

Large positive values mean the significant activation of the memory pattern. Furthermore, we calculated pattern correlation between attractor patterns obtained from different trigger nodes.

LAM converges to various attractor patterns depending on trigger nodes and the value of auto-associative weight *α*. Generally, memory recall (large positive maximum pattern overlap) was observed in the parameter region *α* > −1 (Figure 2d,i,n). When *α* takes large positive values, attractor patterns locally represent one or a few nodes in the graph (Figure 2b,g,right) and patterns for different nodes are uncorrelated to each other (Figure 2c,h,right). These attractors correspond to those obtained by conventional Hopfield-type models. In contrast, when *α* is negative (−1 < *α* < 0), multiple memory patterns are active in attractor states. Quantitatively, the average number of active patterns is maximum at *α* ≈ −1 and decreases as *α* increases (Figure 2e,j,o; see Methods for the definition of active patterns). Especially in *α* ≈ −1, distributions of pattern overlaps represent large communities in graphs (Figure 2b,g,left), and accordingly, pattern correlation between attractor patterns are high within each community (Figure 2c,h,left). When *α* took an intermediate value, LAM exhibited a representation for a mesoscale community in the four-room graph (Figure 2l, right). This result demonstrates that LAM generates mixed representations for communities in the hetero-associative links by partially recalling multiple memory patterns simultaneously in attractor states. Accordingly, representations for nodes within a community are highly correlated, which agrees with experiments^6,7^.

### Theoretical relationship between LAM and graph Laplacian

Next, we analyze the mathematical mechanism behind representations of LAM. We found that the representations are related to graph Laplacian (GL). GL is a matrix defined for a graphical structure and its eigenvectors are used for various applications. One popular application is graph segmentation (also called as community detection) because it has been shown that signs of elements in GL eigenvectors indicates optimal two-fold segmentation of a graph^27,28^ (examples are shown in Figure 3a-c).GL eigenvectors give segmentation in various levels depending on their eigenvalues (small eigenvalue corresponds to the coarse resolution with large communities), thus combinations of multiple eigenvectors enables multi-level segmentation. This property is utilized for image segmentation^27,28^ for example. In another aspect, GL eigenvectors is also used for nonlinear dimensionality reduction^29^, which gives low-dimensional representations for nodes (data points) in which the structure is represented through similarity. As for the connection to neural representations, GL eigenvectors become grid-like code in the homogenous space, and their distortion brought by inhomogeneity fits with experimental observation of grid cells, and predictive spatial representations in hippocampus can be eigendecomposed into GL eigenvectors^10^. See Methods for the definition of GL and a brief review of mathematical properties.

**Figure 3.**
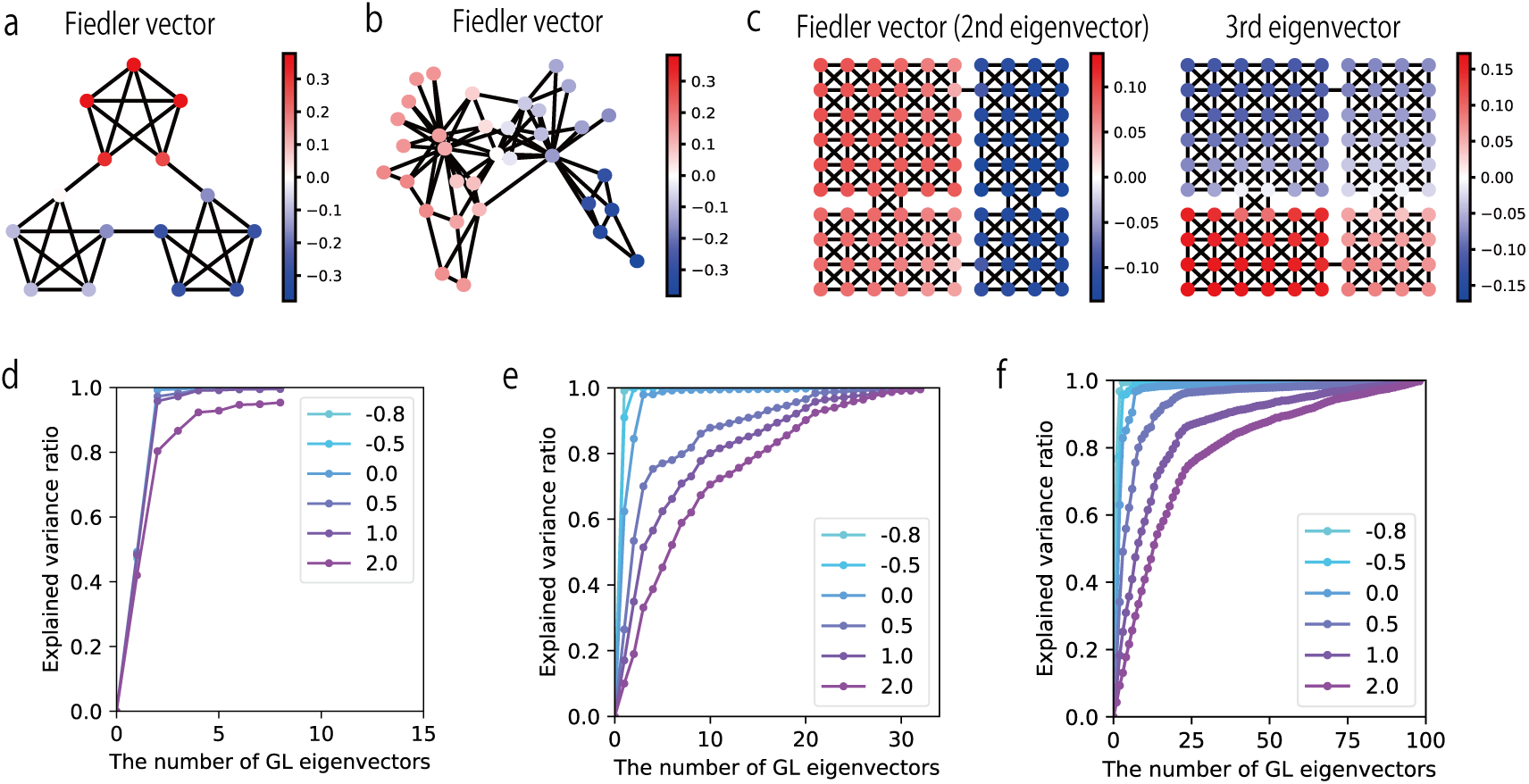
The relationship between Graph Laplacian eigenvectors and LAM. (a-c) GL eigenvectors for example graphs. Fiedler vector is an eigenvector with a second smallest eigenvalue. (a) Fiedler vector for the graph in Schapiro et al. (2013). (b) Fiedler vector for karate-club network. (c) Fiedler vector and the eigenvector with the third smallest eigenvalue for the four-room graph. (d-f) The ratio of variance of pattern overlap vectors explained by various number of graph Laplacian eigenvectors. The color indicates the value of *α*. Values from all attractor pattern triggered by different initial patterns are averaged for each *α*. (d) Results from the graph in Schapiro et al. (2013). (e) Results for karate-club network. (f) Results for the four-room graph.

We performed a formal theoretical analysis to show the relationship between LAM and GL. With symmetric normalization of hetero-associative weights (see Methods for the detail) which gives quantitatively same results as results shown above (Supplementary Figure 1), we can define an energy function of the model as

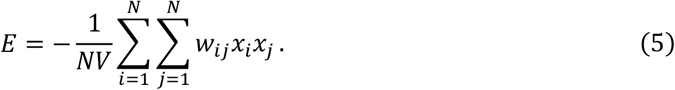

As in the conventional Hopfield model^11^, dynamics of LAM monotonically decreases this energy (Supplementary Figure 2). We consider a vector of pattern overlaps **m** = (*m*^1^, …, *m*^*P*^)^T^ and a vector rescaled by degrees of the graph 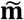, then we can rewrite the energy function as

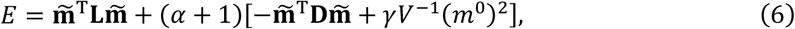

where **L** is GL for the hetero-associative link structure (the state-transition graph) and **D** is a degree matrix, *m*^0^ is the mean activity level in the network. Here we find the minimization of 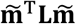 under the constraint of 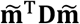, which is the same objective with graph segmentation^27^ and graph-based dimensionality reduction^29^, for which GL eigenvectors give optimal solutions. Therefore, we can expect GL eigenvectors appear in the rescaled pattern overlap vector 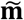 through the energy minimization of LAM. Furthermore, we derived that a GL eigenvector with an eigenvalue *λ*_*k*_ is activated in the pattern overlap vector in the condition *λ*_*k*_ < *α* + 1 (see Methods). Noting that the minimum eigenvalue of GL is always zero and smaller eigenvalue corresponds to coarser graph segmentation, this result indicates that representations of the largest community (the eigenvector with the second smallest eigenvalue, which is called as Fiedler vector) appears in LAM when *α* is slightly higher than −1. As *α* increases, eigenvectors with higher eigenvalues are also activated, thus it is expected that represented communities get smaller. This analysis fits with the results shown in the previous section, especially the similarity between pattern overlaps in *α* ≈ 1 and Fiedler vectors (Figure 2b,g,l and Figure 3a-c). Although this analysis of the energy function depends on symmetricity of synaptic weights, we also derived same activation thresholds of GL eigenvectors through Turing instability analysis for complex networks^35^ which does not require such constraint on connectivity. Therefore, we can expect similar transient dynamical properties also for LAM with asymmetric connections although the existence of attractors is not guaranteed.

Alternatively, we can also interpret the energy minimization above as the combination of two conflicting optimization. First, the minimization of 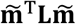 is equivalent to the minimization of differences between pattern overlaps *m*^*μ*^ for two strongly connected cell assemblies^29^. This results in smoothing (or diffusion) on the graph, which leads to non-sparse solutions, observed as mixed representations of multiple cell assemblies. Second, the term 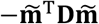 is same as the conventional Hopfield model, which basically leads to the activation of a single memory pattern. Minimization of the mean activity *m*^0^ also helps to create sparse activity patterns. Therefore, the latter part of the energy function acts for the sparsification, that is, reduction of the number of activated memory patterns. In sum, the energy function is composed of two components for smoothness and sparsity, respectively, and the value *α* + 1 determines the trade-off. If *α* < −1, the effect of sparsification vanishes, thus no pattern is preferentially active. The number of activate patterns is maximiazed when *α* is slightly higher than −1 because of the strong smoothing effect. As *α* increases in the region *α* > −1, the number of activated patterns gradually decreases and the model gets close to the conventional Hopfield model. This intuitive interpretation also fits with the actual behavior of the model.

To quantitatively validate the relationship between representations in LAM and GL eigenvectors, we performed linear regression of pattern overlaps of LAM (shown in Figure 2) by various numbers of GL eigenvectors, and calculated variance explained. Eigenvectors were chosen from those with small eigenvalues to those with large eigenvalues. The result shows that pattern overlaps in LAM with low auto-associative weights *α* were mostly explained by small numbers of GL eigenvectors with small eigenvalues, and eigenvectors with large eigenvalues were gradually recruited as *α* increased (Figure 3d-f). This result fits with our theoretical analyses and demonstrates the mathematical mechanism behind representations in LAM.

Based on the relationship with GL, we tested graph-based image segmentation by LAM, which is one of the well-established applications of GL^27^. We assigned a random binary pattern to each pixel and defined hetero-associative links between pixels based on spatial proximity and similarity of RGB values similarly to the previous study^27^. LAM successfully extracted large segments corresponding to a GL eigenvector when the auto-associative weight *α* was close to −1 (Supplementary Figure 3). When *α* was increased, LAM extracted relatively small segments (Supplementary Figure 3). These results show that LAM is also applicable to non-ideal graphs constructed from real-world data.

### Finding subgoals by graph-based representations and novelty detection

One of important applications of GL eigenvectors is to find appropriate subgoals for hierarchical reinforcement learning^8,9^. In this framework, a set of primitive actions (called as options) are optimized through learning to reach subgoals. Desirable subgoals are “bottlenecks” shared by many trajectories on the state-transition graph. GL eigenvectors have been used to find bottlenecks through graph segmentation. We tested whether representations in LAM can also be used for subgoal finding by comparing results between LAM and GL.

To identify bottlenecks, we calculated “novelty index” of each node, which measures the expected changes of representations caused by the movements from a node to surrounding nodes (see Methods for the mathematical definition). In hierarchical reinforcement learning, subgoals are treated as pseudo-reward for agents. It is biologically natural to treat novelty as pseudo-reward because dopamine cells are activated by not only reward but also novelty^36^. With GL, we constructed low-dimensional representations of nodes from GL eigenvectors with low eigenvalues (Laplacian eigenmap)^29^. On the other hand, with LAM, activity patterns in attractor states are directly used as representations.

With GL, the novelty index successfully detected nodes located at the bottlenecks (doorways) between distinct compartments (Figure 4a). Furthermore, the sensitivity of bottleneck detection was controlled by the dimension of representation vectors: the use of a single eigenvector with low eigenvalue (Fiedler vector) extracted the narrowest bottlenecks, and higher dimensional representations enabled the detection of other bottlenecks. We obtained equally good performance by LAM (Figure 4b). In LAM, auto-associative weight *α* regulates the number of active GL eigenvectors in representations, hence it changes the sensitivity of bottleneck detection. This result demonstrates that novelty detection in LAM enables multi-resolution subgoal finding comparable to GL eigenvectors. The idea of using novelty has been suggested in the literature of hierarchical reinforcement learning^37^ but LAM provides a more biologically plausible mechanism based on a neural network.

**Figure 4.**
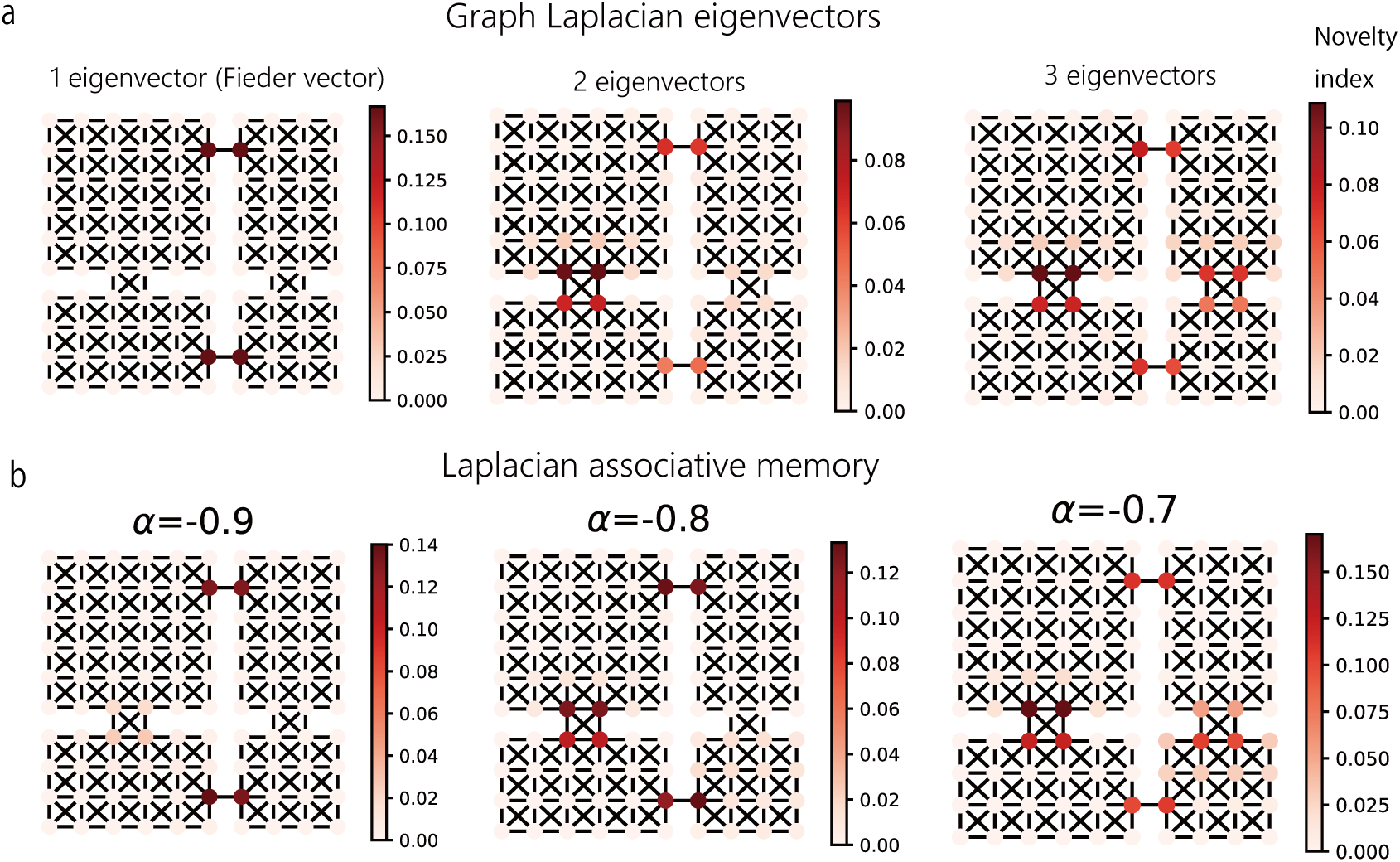
Subgoal finding by novelty detection with representations in LAM. (a) Subgoal finding using low-dimensional (1, 2, or 3) representations constructed from graph Laplacian eigenvectors. The color indicates the novelty index for each node. (b) Subgoal finding using representations in LAM obtained by different values of *α*.

### Chunked sequential activities in asymmetric LAM

So far, we have analyzed attractor patterns in LAM with symmetric links. Next, we show dynamic properties of asymmetric LAM. We constructed asymmetric LAM with a ring-shape graph in which link weights were slightly stronger in one direction than in the opposite direction (Figure 5a). We simulated the neural activity while continuously changing the value of the auto-associative weight *α* (Figure 5b, top). The network showed a sequential activity in which embedded memory patterns are consecutively retrieved at a variable speed (Figure 5b, bottom). Rapid state transitions selectively occurred when the value of *α* became negative and close to −1 (Figure 5d), at which the distribution of pattern overlaps was maximally expanded (Figure 5c). These results indicate that negative auto-associative weights in asymmetric LAM not only generate macroscopic representations for large communities but also increases the sensitivity to asymmetricity in synaptic weights and facilitates sequential transitions across memories.

**Figure 5.**
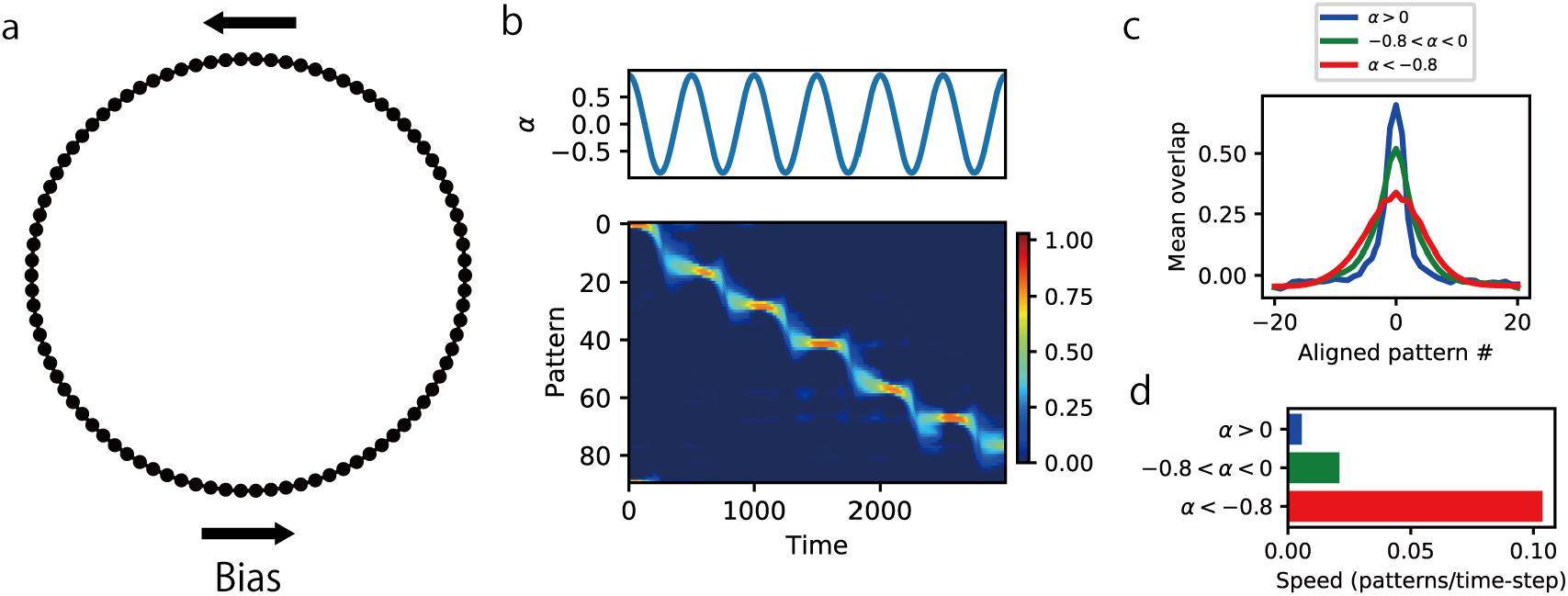
Parameter-dependent sequential activities in asymmetric LAM. (a) A ring structure of hetero-associative links for the simulation of asymmetric LAM. Hetero-associative weights were biased towards one direction. (b) The time course of *α* (top) and pattern overlaps (bottom) in the simulation of asymmetric LAM. Negative pattern overlaps were truncated to zero. (c) Peak-aligned mean pattern overlap distributions in different ranges of *α*. (d) Mean speed of the peak shift of the pattern overlap distribution in different ranges of *α*.

Motivated by this dynamic property and the relationship between LAM and GL, we examined whether the sequential activities in asymmetric LAM are chunked according to the communities in hetero-associative links. For simulations, we specifically focused on hippocampal theta sequences in which chunking was experimentally observed^32^. We assumed a virtual animal running on a ring-shape track, and we modeled the hippocampus of the animal by asymmetric LAM with a ring-shape hetero-associative link structure (Figure 6a). We simulated neural activities of LAM with the fixed parameter *α* = −0.9 while regularly stimulating a cell assembly encoding the current location of the animal. In the simulation, we obtained rhythmic sequential activities along the ring (Figure 6b), which are a simplified model of theta sequences.

**Figure 6.**
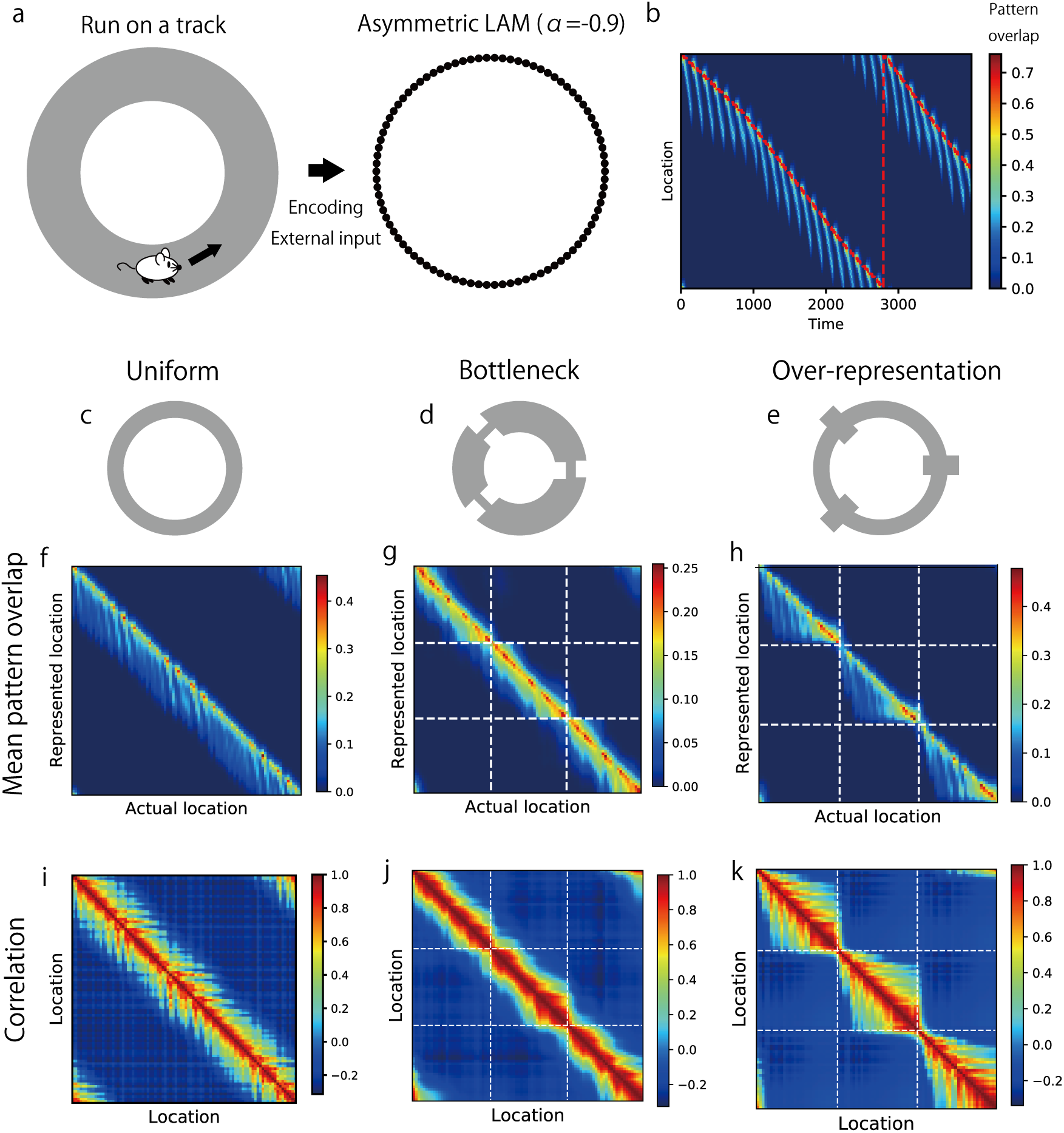
Chunked sequential activities in asymmetric LAM. (a) Schematics for the simulation setting. We assumed a virtual animal running around a ring track, and regularly stimulated patterns in asymmetric LAM corresponding to the current location. (b) Patten overlaps of simulated neural activities in the uniform model. The red dotted line indicates the actual location of the virtual animal. (c-e) Schematics of the hetero-associative link structure. A uniform ring (c), a ring chunked with local bottlenecks (d), and a ring chunked with local over-representations (f). Details are shown in supplementary figure 4. (f-h) Mean pattern overlaps (the strength of representations of locations) at each actual location of a virtual animal in the simulation of a uniform model (Ff), a bottleneck model (g), and over-representation model (h). We truncated negative pattern overlaps to zero in these figures. (i-k) Correlations between mean pattern overlaps at different locations in the simulation of a uniform model (i), a bottleneck model (j), and over-representation model (k). White dotted lines in g, h, j, k indicate locations of chunk borders (bottlenecks or over-representations).

In this model, we tested three hetero-associative link structures (see Supplementary Figure 4 for the detail of structures). In a uniform ring structure without chunks (Figure 6c) sequential activities propagated homogenously (Figure 6f, i). When the structure has local bottlenecks (Figure 6d), sequential propagation was constrained at the bottlenecks (Figure 6g,j). This result is analogous to that of symmetric LAM with the four-room graph (Figure 2k). Finally, when the structure is chunked by local over-representations (Figure 6e) which were implemented as densely connected nodes, sequential propagation was also chunked at over-represented locations and representations strongly correlated within chunks (Figure 6h,k). Although both the bottleneck model and the over-representation model exhibited chunking effects, the over-representation model is particularly consistent with the experimental observation because theta sequences are segmented at salient landmarks and rewards^32^ which are over-represented by hippocampal place cells^38,39^. These results demonstrates that LAM provides a unified mechanism for graph-based representations^6,7^ and chunking of sequential activities^32^.

## Discussion

In this paper, we proposed Laplacian associative memory (LAM), an extension of the Hopfield-type network models to compute community structures in hetero-associative links. While structural segmentation has been attempted by hierarchical networks with different time constants^40^, our model provides a novel framework for multiscale information processing in a single network and accounts for experimentally observed graph-based representations^6,7^. Furthermore, we showed that LAM with asymmetricity can generate chunked sequential activities which reproduced experimentally observed chunking of theta sequences^32^. Notably, a model parameter crucial for segmentation can be interpreted as the strength ratio of local (assembly-specific) and global (non-specific) inhibitory feedback. This interpretation offers a novel insight into the computational roles of inhomogeneous neuromodulations of interneuron circuits, which is a focus of recent experimental studies^30,31^.

We interpreted the model parameter *α* that controls the scale of the representation as the balance between global and local inhibition. In the light of recent experimental evidence, we further speculate the specific circuit mechanisms to regulate the parameter *α*. In visual cortex, somatostatin-expressing (SOM) interneurons and palvalbumin-expressing (PV) interneurons are considered to serve for anatomically global and local inhibitory feedback, respectively^41^. Furthermore, the cholinergic activation of vasoactive intestinal peptide-expressing (VIP) interneurons selectively inhibits SOM interneurons, which changes the balance between SOM and PV interneurons^30,31^. Then, it seems reasonable to hypothesize that acetylcholine elicits a local-inhibition-dominant (PV-dominant) state in cortical circuits and hence generates the macroscopic representations corresponding to large communities. This hypothesis is consistent with the computational model of cortical inference in which acetylcholine signals uncertainty^42^ because acetylcholine creates mixed and ambiguous representations over many states. However, our model offers a novel prediction that uncertainty of the inference (expansion of probabilistic distributions) is constrained by communities in the graph structure behind experience. Our hypothesis also predicts that the malfunctioning of PV interneurons results in the deficit of processing uncertainty and macroscopic information, and difficulty of transition between states. This may be consistent with the symptom and the likely cause of schizophrenia^43–45^. These possibilities should be pursued by more specific and detailed modeling and fitting to experimental data.

We used graph-based representations in LAM for subgoal finding in HRL^8,9^. Another way to perform reinforcement learning with graph-based representations is to use successor representation^46^. Successor representation gives prediction of near-future state occupancy from current states that is useful for value estimation, and has been shown to be consistent with many experimental findings in hippocampal information representations^10^ including representational similarity within community structures shown in this study. Our model shares similar properties with successor representation as its eigendecomposition yields GL eigenvectors^10^ and modifying timescale of representation results in changing eigenvalues. Therefore, we expect some theoretical connection between successor representation and our model. However, we note that our model directly exhibits the Fiedler vector in *α* ≈ −1, but successor representation cannot give such representation in any parameter setting (see Methods). Therefore, in the current form, there is a little quantitative difference between our model and successor representation. Which model better explains experimental observations and whether our model (with some modification) exhibits successor representation are interesting open problems.

In asymmetric LAM, we found that negative auto-associative weights facilitate sequential transitions across memory patterns. Previously, we found that negative auto-association significantly increases the sensitivity of correlated attractors to external perturbation^23^. We speculate that changes in the propagation speed presented here depends on a similar mechanism. If the auto-association is strongly positive, attractors are stable and invulnerable to directional biases in link weights. However, as *α* gets close to −1, the attractors are gradually instabilized and become sensitive to weight biases and external perturbations. This property suggests that macroscopic representations are dynamic in the brain, and are unlikely to serve for robust working memory in the brain as conventional attractor networks^20,24,25^.

We found that both local bottlenecks and over-representations induce the chunking of sequential activities in asymmetric LAM. The over-representation model is particularly interesting because it accounts for the role of salient landmarks and rewards which are over-represented by place cells^38,39^. We may be able to apply a ring-shape structure with two over-representations for modeling typical experiments in which animals run back and forth on a 1-D track to get rewards at both ends, considering that many place cells are direction-selective in such experimental setting^47^. In contrast, to our knowledge, how bottlenecks affect hippocampal sequential activities has not been tested experimentally. An adequate design of bottlenecks does not seem to be trivial in spatial navigation tasks because animals may recognize spatial bottlenecks as salient landmarks which would be over-represented in the brain. A proper design of the task structure requires a careful control of saliency of each state.

The simple model with the asymmetric LAM produced sequential activities similar to chunked hippocampal theta sequences^32^ (Figure 6). However, the hippocampal circuits generate more complex oscillatory dynamics, which is also likely to contribute to segmentation. For instance, in hippocampal replays of spatial trajectories, a boundary of chunks (a bifurcating point) in the spatial structure is locked to troughs of LFP power in concatenated sharp wave ripples^33^. Furthermore, hippocampal circuits repeat convergence to and divergence from discrete attractors every gamma cycle during sharp wave ripples^16^. Our simplified model cannot address the relationship between such complex oscillatory dynamics and segmentation. A detailed network model involving realistic spiking neurons and inhibitory circuits is necessary for studying such the relationship.

Previously, processing hierarchical knowledge in associative memory models was implemented by embedding artificially correlated memory patterns^48^. This model successfully reproduced the dynamics of hierarchical information processing in the temporal visual cortex^49,50^. The relationship between our model with hetero-associative links and the previous model with correlated memory patterns is currently unclear, and worth exploring. If similar graphical computation is possible with correlated memory patterns, the brain may perform graphical computation based on not only temporal association (hetero-associative links in our model) but semantic similarity between items (correlation between memory patterns). However, we emphasize that our finding that associative memory networks can autonomously compute mathematically well-defined communities in complex graphs is previously not known, because previous models have tested only simple structures and negative auto-association has not been considered.

The mechanism proposed in this paper will give a novel method to solve an arbitrary eigenvalue problem by using associative memory models. In the present model, we constructed hetero-associative weights from normalized adjacency matrices of graphs. However, the proposed dynamical mechanism to solve eigenvalue problem is generic and does not depend on this specific condition. For example, if we employ covariance matrix between encoded variables for a hetero-associative weight matrix, a network is theoretically expected to perform principal component analysis. Because eigenvalue problems ubiquitously appear in applied mathematics and machine learning, other computational methods may also be mapped to brain functions through the similar mechanism. Our model suggests much more powerful computing ability of associative memory models than previously thought and may provide a bridge across artificial intelligence and brain science.

## Acknowledgements

We are grateful to H. Shiwaku for helpful discussion. We also thank OIST Scientific Computation and Data Analysis section for the technical support for scientific computing. This work was partially supported by Kakenhi nos. 18H05213 and 19H04994 from the MEXT (Ministry of Education, Culture, Sports, Science and Technology, Japan).

## Author contributions

T. H. and T. F. conceived the project and wrote the manuscript. T. H. mathematically designed the model, and performed simulations and analyses.

## Competing financial interests

The authors declare no competing interests.

## Methods

### Definition and mathematical properties of graph Laplacian

Let us assume a symmetric graph which has an adjacency matrix **A** whose element *A*_*ij*_ denotes the existence of an edge with 0 and 1 (unweighted graphs) or the weight of the edge (weighted graphs) between the node *i* and the node *j*. We also define a degree matrix **D**, in which diagonal elements are degrees (the number of edges connected to each node) *d*_*i*_ = ∑_*j*_*A*_*ij*_ and other elements are zero. Graph Laplacian is a matrix defined as **L** = **D** − **A**. There are two ways of normalization: a symmetric one 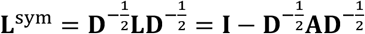 and an asymmetric one **L**^asym^ = **D**^−1^**L** = **I** − **D**^−1^**A** (**I** is an identity matrix). These two matrices have qualitatively same properties^28^.

An important characteristic of graph Laplacian matrix is that its eigenvectors give optimal graph segmentation. Here optimality is defined by the min-cut criterion that prefers a two-fold division of a graph obtained by cutting the minimum number of edges. It has been proven that min-cut graph segmentation can be performed by solving the generalized eigenvalue problem **Ly** = *λ***Dy**, or equivalently, eigenvectors of normalized graph Laplacian **L**^sym^ and **L**^asym27,28^. A sign of each element in the eigenvector **y** indicate a segment that each node should be assigned, and multiple eigenvectors correspond to the two-fold segmentation in various levels, depending on their eigenvalues. The eigenvector with the second smallest eigenvalue (Fiedler vector) is regarded as the best non-trivial solution which corresponds to the largest community structure (which achieves minimum cut) in the graph. Eigenvectors with larger eigenvalues are suboptimal solutions perpendicular to other eigenvectors, and tend to give subdivision of large communities into subclusters.

Another useful interpretation of graph Laplacian eigenvectors is low-dimensional representations for nodes in the graph which is called as Laplacian eigenmap^29^. The generalized eigenvalue problem **Ly** = *λ***Dy** gives perpendicular solutions for min 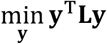 subject to **y**^T^**Dy** = 1 and the eigenvalue indicates the minimized value. Because of the relationship

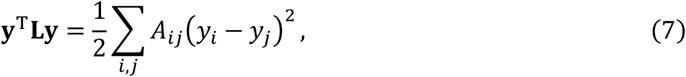

minimization of **y**^T^**Ly** can be regarded as assigning values *y*_*i*_ to nodes such that strongly connected nodes are represented by close values. In this sense, low-dimensional representations constructed from graph Laplacian eigenvectors with low eigenvalues captures the graph structure through their similarity, which is the appropriate property for nonlinear dimensionality reduction.

### Construction of hetero-associative weights

In this study, we hypothesize that hetero-associative weight matrix 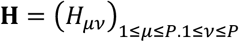 is constructed as 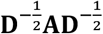 (symmetric normalization) or **D**^−1^**A** (asy m metric norma lization) where **A** and **D** are the adjacency matrix and the degree matrix of the graph, respectively. As in graph Laplacian, two normalization yields qualitatively same results. However, the symmetric normalization model enables us formal theoretical analyses and the asymmetric normalization model gives biologically plausible interpretation of the model.

Asymmetrically normalized weights directly correspond to the transition probability matrix for random walk on the graph^28^, thus they can be naturally learned in sequential experiences through Hebbian learning at the state transition^22,25^. With some additional assumptions, we can specify the learning rule for weights of asymmetric normalization model. We assume that an agent experiences a sequence of discrete states *s*[*t*], and the neural activity *r*_*i*_[*t*] exhibits a binary pattern 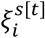 corresponding to the current state. We consider an activity-dependent learning rule:

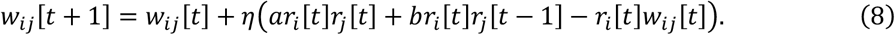

In a stable state, E [*w*_*ij*_[*t* + 1] − *w*_*ij*_[*t*]] = 0 and E [*w*_*ij*_[*t*]]can be regarded as a constant *w*_*ij*_. Then,

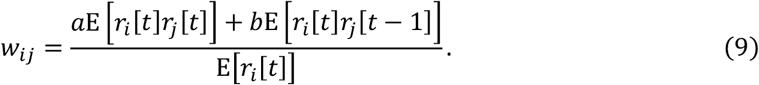

Using 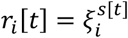,

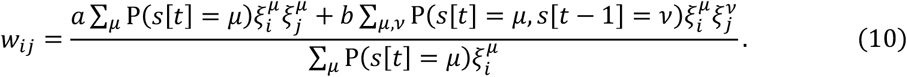

If patterns 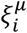 are sparse and overlaps between patterns are negligible (each neuron participates a single pattern at most),

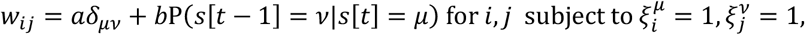

and *w*_*ij*_ = 0 if neuron *i* or *j* does not belong to any pattern (assuming initial weights are zero). Finally, with 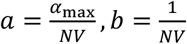 and P(*s*[*t* − 1] = *ν*|*s*[*t*] = *μ*) = *H*_*μν*_, we obtain

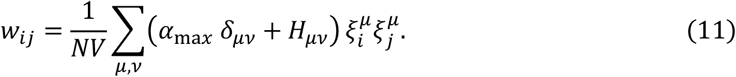

In a random walk on a state-transition graph, *H*_*μν*_ = P(*s*[*t* − 1] = *ν*|*s*[*t*] = *μ*) corresponds to an element in the normalized adjacency matrix **D**^−1^**A**.

### Decomposition of excitatory and inhibitory synaptic weights

With the asymmetric normalization model, synaptic weights can be decomposed as

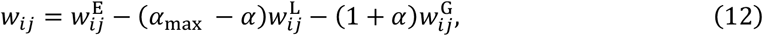

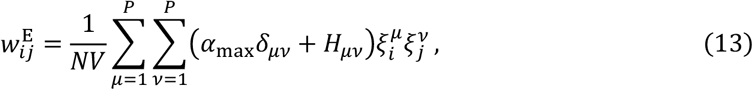

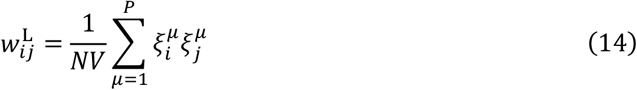

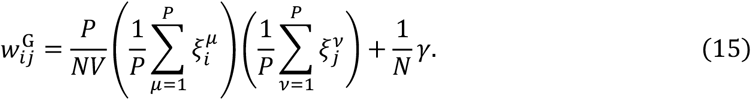

Here, we used the constraint 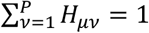 and approximated 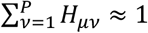. All decomposed weights are always non-negative. The component 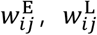, and 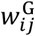 are excitatory connections (which reflects the structure of cell assemblies), assembly-specific local inhibition, and non-selective global inhibition, respectively.

### Analysis of the energy function of LAM

Here we consider symmetric normalization model in which hetero-associative links are constructed as 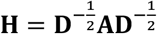. As in the main text, the energy function is

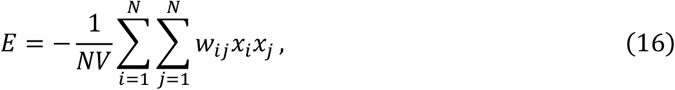

We define a pattern overlap

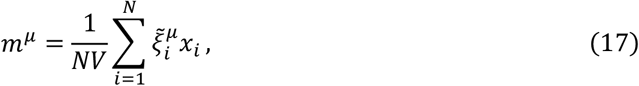

a pattern overlap vector **m** = (*m*^1^, …, *m*^*P*^)^T^, and the mean activity 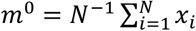. Then, we can rewrite the energy function as

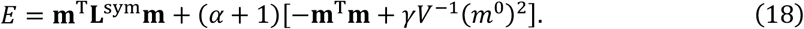

The matrix **L**^sym^ = **I** − **H** is symmetric normalized graph Laplacian^27^ if we regard the hetero-associative weight matrix as the normalized adjacency matrix. By rescaling **m** by the degree matrix of the hetero-associative links **D** as 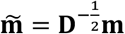, we further obtain

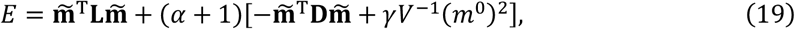

where **L** is unnormalized graph Laplacian.

To see the relationship with graph Laplacian eigenvectors more quantitatively, we expand the overlap vector **m** by a linear combination of eigenvectors of symmetric normalized graph Laplacian **ϕ**_*k*_ (with corresponding eigenvalues *λ*_*k*_) as

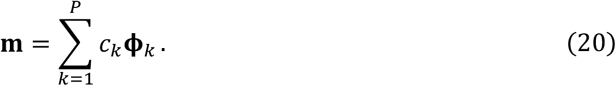

Then, the energy function can be written as

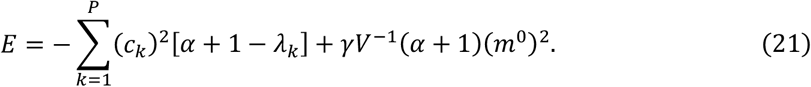

If *γ* = 0, the minimization of this energy requires *c*_*k*_ ≠ 0 if *λ*_*k*_ < *α* + 1, which gives the approximate threshold of the activation of an eigenvector **ϕ**_*k*_ in the representation (note that the actual threshold can be shifted because of *γ* > 0).

### Turing instability analysis of LAM

For the analysis, we first replace the step function Θ(*x*) in Eq. (1) by a differentiable monotonically increasing function f(*∈x*) that converges to Θ(*x*) in the limit of *∈* → ∞ (e.g. a logistic function f(*∈x*) = (1 + exp(−*∈x*))^−1^). As in the main text, we define pattern overlaps as

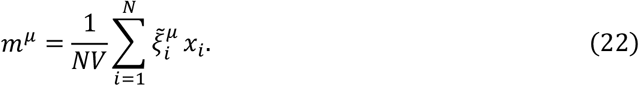

From Eq. (1)(2), dynamics of overlaps can be obtained as

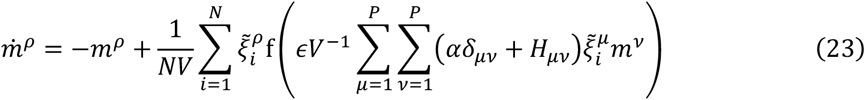

Next, with vectors **m** = (*m*^1^, …, *m*^*P*^)^T^, 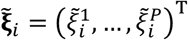, and the hetero-associative weight matrix 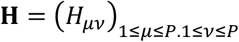, we obtain a vector representation as

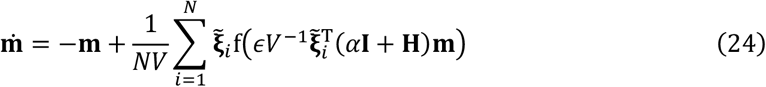

where **I** is an identity matrix. When *N* and *P* are sufficiently large and memory patterns are random, **m** = **0** is an equilibrium point for this dynamical equation. Furthermore, in that condition, the matrix 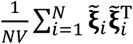 becomes a correlation matrix for random memory patterns, which can be approximated by an identity matrix. Thus, we obtain the following equation by linearizing f(*x*) around **m** = **0**:

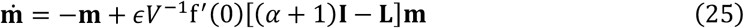

Here, we defined **L** = **I** − **H** (this is either symmetric or asymmetric normalized graph Laplacian). Finally, we expand **m** with eigenvectors **ϕ**_*n*_ (*n* = 1, …, *P*) of the matrix **L** as

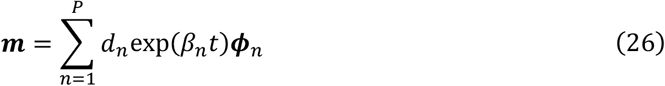

Substituting this into the linearized equation yields

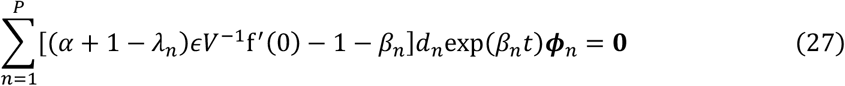

where *λ*_*n*_ is an eigenvalue for ***ϕ***_*n*_. This equation has non-trivial solutions (*d*_*n*_ ≠ 0) only if *β*_*n*_ = (*α* + 1 − *λ*_*n*_)*∈V*^−1^f′(0) − 1, which gives exponential growth rates along each eigenvector around **m** = **0**. If there exists a positive growth rate, the network becomes instable along the corresponding eigenvectors; otherwise the network is stabilized at **m** = **0**. In the limit of *∈* → ∞ (f(*∈x*) → Θ(*x*)), the sign of *β*_*n*_ is solely determined by the sign of *α* + 1 − *λ*_*n*_. This result suggests that the overlap vector **m** is activated (instabilized) along the *k*-th eigenvector of the graph Laplacian matrix **L** if *α* > *λ*_*k*_ − 1 (*λ*_*k*_ is the eigenvalue for the *k*-th eigenvector).

### Simulations of the network model

In numerical simulations, we used decomposed asymmetric normalization model unless specified otherwise. We first initialized activities by one of memory patterns 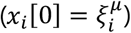 and updated activities by a discretized version of Eq. (1):

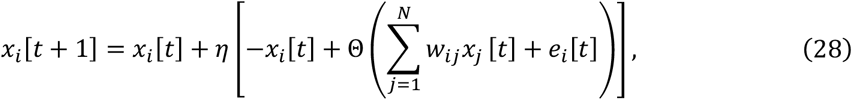

where *e*_*i*_ [*t*] is an external input applied in simulations in Figure 6. We used *η* = 0.01 for simulations with symmetric graphs (Figure 2, 3, 4 and Supplementary Figure 1, 2, 3) and *η* = 0.1 for simulations of sequential activities (Figure 5 and 6). The number of neurons were *N* = 30000 for image segmentation tasks, and *N* = 10000 for all other simulations. The additional inhibition parameter was *γ* = 0.6 for image segmentation and simulations of sequential activities, and *γ* = 0.3 for all other simulations. Sparsity *p* was 0.1 (approximately 10% of neurons are active in each pattern) throughout the study.

Attractor patterns of the network model with symmetric graphs were obtained by simulations of 3,000 time-steps. Simulations of sequential activities in Figure 5 was performed for 10,000 time-steps. Simulations of sequential activities in Figure 6 were performed for 30,000 time-steps and repeated three times using different random seeds for each setting. We averaged mean pattern overlaps at each location and correlations between mean pattern overlaps over those three trials. We truncated negative mean pattern overlaps to zero in this calculation.

We counted the number of active patterns by counting the number of *μ* that satisfies *m*^*μ*^ > 0.05 and 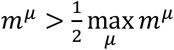.

### Settings for image segmentation

For image segmentation tasks, we took images from pixabay (https://pixabay.com/). We trimmed and down-sampled images so that it contains 1000 - 1500 pixels (e.g. *P* ≈ 1000). We note that images shown in figures are ones before down-sampling. We constructed link weights by the same way with Shi & Marik (2000)^27^:

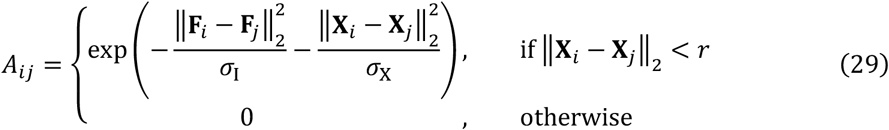

where vectors **F**_*i*_ and **X**_*i*_ denotes the RGB value (normalized between 0 and 1) and the spatial location of pixel *i*, respectively. Parameters were *σ*_I_ = 0.1, *σ*_X_ = 4, *r* = 5. After setting values, we performed asymmetric normalization of weights to get hetero-associative weights.

### Definition of the novelty index for subgoal finding

First, we calculate a representation of each node by either constructing low-dimensional vectors from graph Laplacian eigenvectors (Laplacian eigenmap)^29^ or simulating an attractor pattern of LAM triggered by a memory pattern of each node 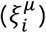. We define similarity between representations of two nodes as *s*(*μ*, *ν*) by either cosine similarity between two representations of Laplacian eigenmap, or correlation between attractor patterns of LAM. The novelty index of a node *μ* is defined as

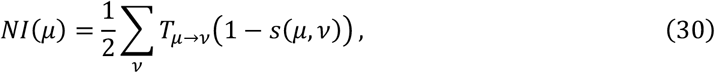

where *T*_*μ*→*ν*_ denotes the transition probability from *μ* to *ν* in random walk on the graph (which is equivalent with elements in **D**^−1^**A**). The novelty index *NN*(*μ*) spans from 0 to 1 and indicates the expected change of information representations that an agent experience in the transition from the node *μ*.

### Asymmetric Laplacian associative memory and the model of a virtual animal

To construct asymmetric hetero-associative weights, we converted symmetric graphs into mutually connected asymmetric weighted graphs. We set the weight of asymmetric links in the biased direction (including diagonal connections) to 110, and the weight for opposite directions to 90. All links horizontal to the biased direction (radial connections) was 100. After constructing adjacency matrices, we performed asymmetric normalization as in symmetric graphs.

In the simulation in Figure 6, we represented the current location of the virtual animal on the track by a continuous value *z*[*t*] ranging from 0 to 90 which corresponds to 90 nodes in the uniform ring-shape graph (uniform model). The velocity *z*[*t* + 1] − *z*[*t*] was constant but we performed resampling of the velocity from a range [0.02, 0.04] at the timings determined by Poisson process (the mean interval was 1000 time-steps). We determined the index of stimulated pattern by truncating *z*[*t*] to an integer. We applied stimulation to the network every 150 time-steps (the uniform model and the over-representation model) or 200 time-steps (the bottleneck model). The amplitude and the length of stimulation was 0.3 and 50 time-steps, respectively.

In the bottleneck model and the over-representation model, we connected additional nodes at the side of the uniform ring-shape graph (as shown in Supplementary Figure 4). We did not stimulate patterns corresponding to additional nodes. We calculated the pattern overlap for each location by averaging over nodes in the central ring and additional nodes at the same location.

### Eigenvalues of successor representation

Successor representation is defined for a pair of states *s* and *s*′ as

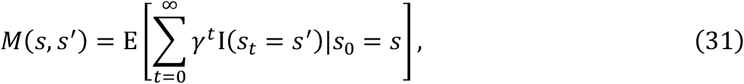

where *γ* indicates a discount factor. We consider a matrix of successor representation **M**, a transition probability matrix **T**, and **L**^asym^ = **I** − **T** is an asymmetric normalized graph Laplacian. Eigenvectors and eigenvalues of **L**^asym^ are defined as **ϕ**_*i*_ and 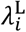, respectively. Then, they satisfy

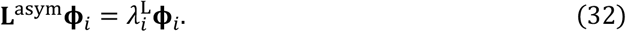

Using the relationship **M** = (**I** − *γ***T**)^−110,46^, we can rewrite this equation with a successor representation matrix:

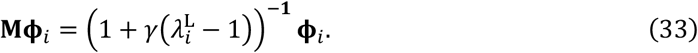

Therefore, eigenvectors of the successor representation matrix are equivalent to those of graph Laplacian, and eigenvalues are 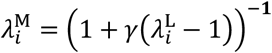. This relationship becomes 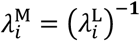 in the limit of *γ* → 1, and 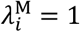 in the limit of *γ* → 0. Thus, although the contribution of Fiedler vector increases as *γ* goes to 1, it is impossible to have 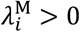 for only Fiedler vector if the graph has a sufficiently complex structure and 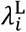 is continuously distributed.

## Notes

### Competing Interest Statement

The authors have declared no competing interest.

