## Supplementary figures for "Associative memory networks for graph-based abstraction"

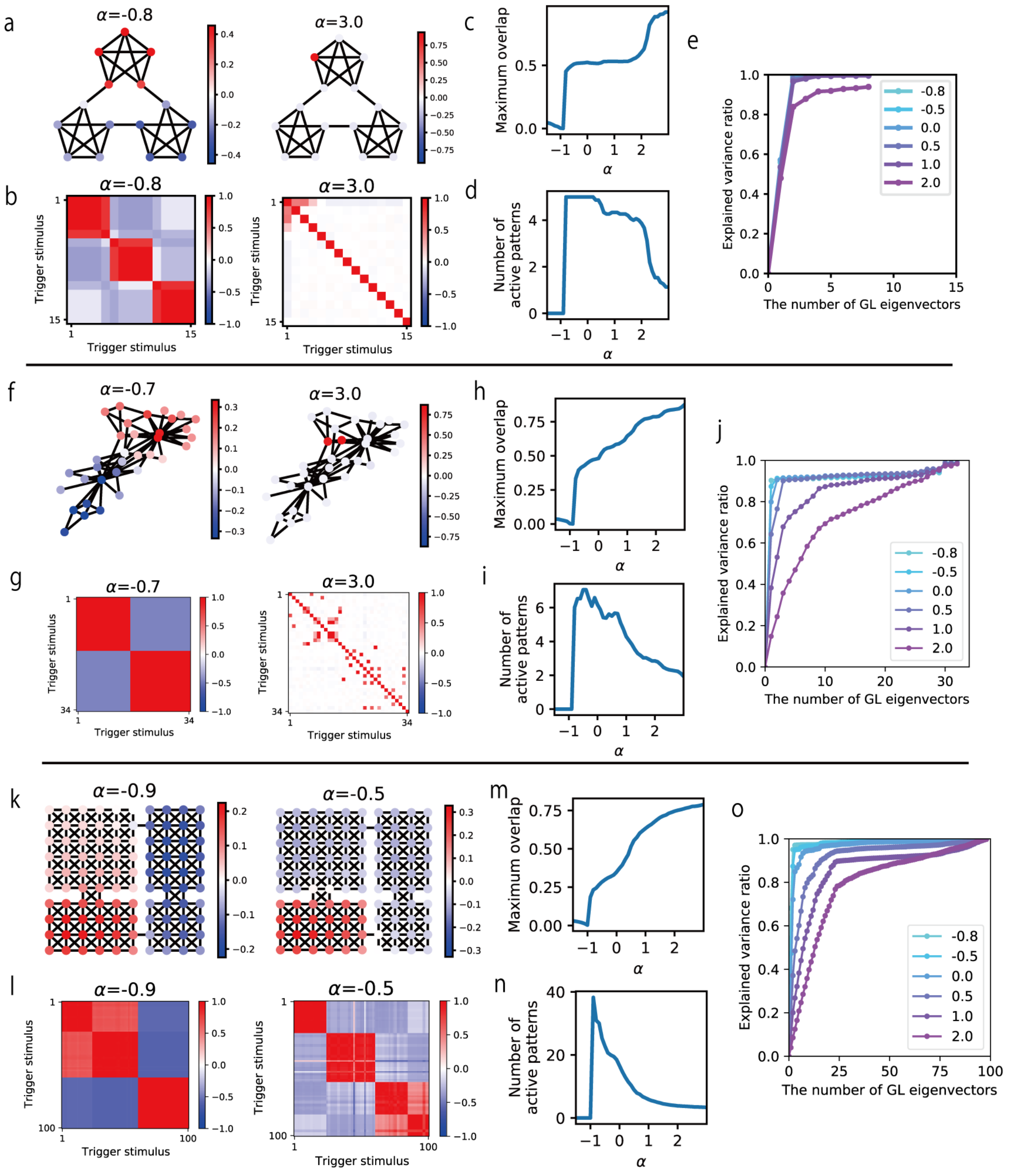


**Supplementary Figure 1** LAM (symmetric normalization model) extracts multi-scale representations for community structures. (a) Pattern overlaps of example attractor patterns. (b) Correlation matrices between attractor patterns triggered by different initial patterns obtained. (c) Maximum pattern overlaps obtained by various values of $\alpha$. (d) Numbers of active patterns obtained by various values of $\alpha$. (e) The ratio of variance of overlap distributions explained by various number of graph Laplacian eigenvectors. The color indicates the value of $\alpha$. We note that, in figures c-e, values from all attractor pattern triggered by different initial patterns are averaged for each $\alpha$. (g-j) Results for Karate-club network. (k-o) Results for compartmentalized rooms.

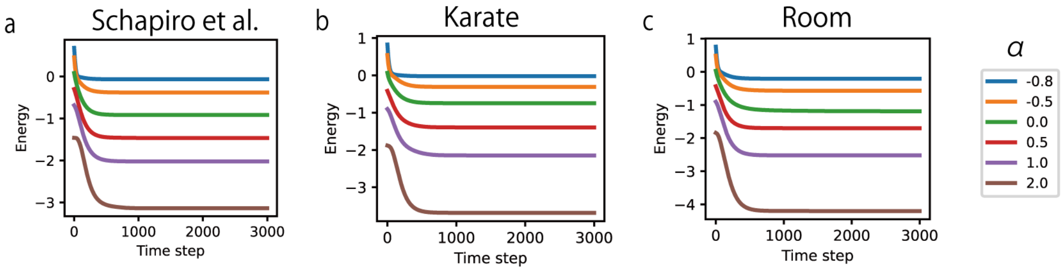


**Supplementary Figure 2** The change of the energy function in the simulation of symmetric normalization model. (a) The graph used in Schapiro et al. (2013). (b)Karate club network. (c) The four-room graph.

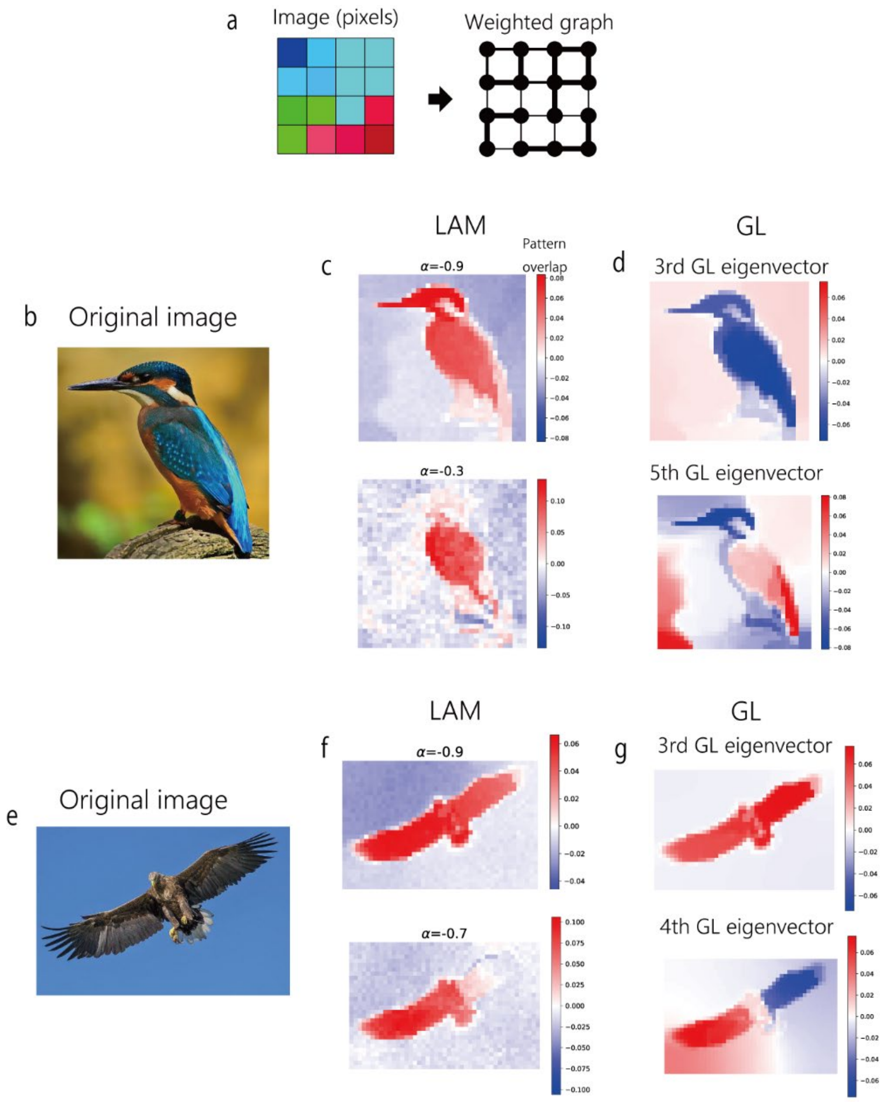


**Supplementary Figure 3** Image segmentation by LAM. (a) Conversion of images into a weighted graph. We regarded each pixel as a node and determined link weights by spatial proximity and similarity of RGB values. (b,e) Original hi-resolution images used for the segmentation task. We used down-sampled images for the construction of graphs. (c,f) Pattern overlaps obtained after the simulation of LAM with different values of $\alpha$. (d,g) Representative GL eigenvectors corresponding to segments obtained by LAM.

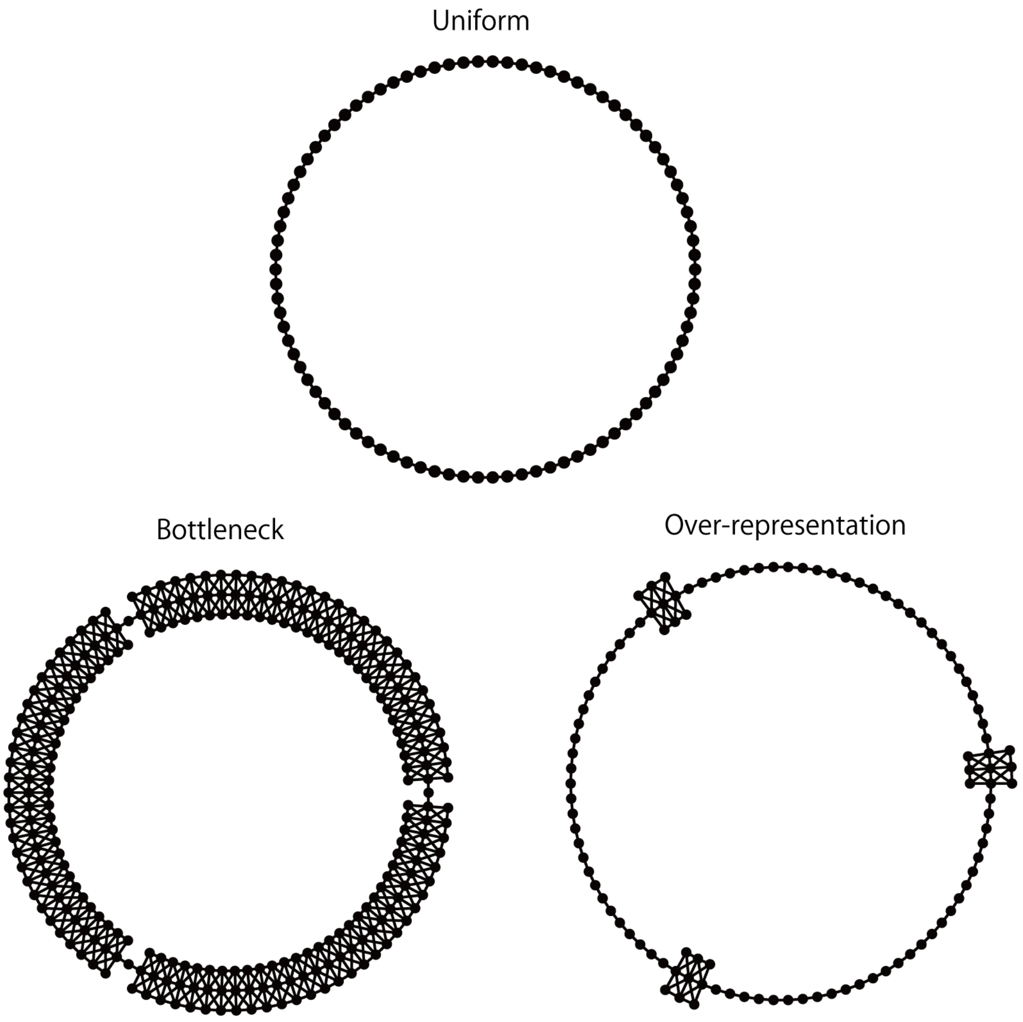


**Supplementary Figure 4** Chunked structures of hetero-associative links used for asymmetric LAM.
